# Opposing roles of p38α-mediated phosphorylation and arginine methylation in driving TDP-43 proteinopathy

**DOI:** 10.1101/2021.08.04.455154

**Authors:** Mari Aikio, Heike J. Wobst, Hana M. Odeh, Bo Lim Lee, Bradley Class, Thomas A. Ollerhead, Korrie L. Mack, Alice F. Ford, Edward M. Barbieri, Ryan R. Cupo, Lauren E. Drake, Nicholas Castello, Ashmita Baral, John Dunlop, Aaron D. Gitler, Ashkan Javaherian, Steven Finkbeiner, Dean G. Brown, Stephen J. Moss, Nicholas J. Brandon, James Shorter

## Abstract

Amyotrophic lateral sclerosis (ALS) is a fatal neurodegenerative disorder typically characterized by insoluble inclusions of hyperphosphorylated TDP-43. The mechanisms underlying toxic TDP-43 accumulation are not understood. Persistent activation of p38 mitogen-activated protein kinase (MAPK) is implicated in ALS. However, it is unclear how p38 MAPK affects TDP-43 proteinopathy. Here, we demonstrate that inhibition of p38α MAPK reduces pathological TDP-43 phosphorylation, aggregation, cytoplasmic mislocalization, and neurotoxicity. We establish that p38α MAPK phosphorylates TDP-43 at pathological serine 409/410 (S409/S410) and serine 292 (S292), which reduces TDP-43 liquid-liquid phase separation (LLPS) but allows pathological TDP-43 aggregation. Moreover, we show that protein arginine methyltransferase 1 methylates TDP-43 at R293. Importantly, S292 phosphorylation reduces R293 methylation, and R293 methylation reduces S409/S410 phosphorylation. R293 methylation permits TDP-43 LLPS and reduces pathological TDP-43 aggregation. Thus, strategies to reduce p38α-mediated TDP-43 phosphorylation and promote R293 methylation could have therapeutic utility for ALS and related TDP-43 proteinopathies.

## Introduction

Amyotrophic lateral sclerosis (ALS) is a fatal disorder caused by degeneration of motor neurons (Wobst et al., 2020). While most ALS cases (∼90–95%) are considered sporadic with unknown etiology (sALS), ∼5–10% of cases are familial in nature (fALS), exhibiting a dominant pattern of inheritance (Rowland and Shneider, 2001; Valdmanis and Rouleau, 2008). ALS is linked to mutations in more than 25 genes, with the C9ORF72 hexanucleotide repeat expansion and mutations in the copper–zinc superoxide dismutase (SOD1) being the most common genetic causes (Nguyen et al., 2018). Although mutations in Transactive Response DNA binding protein 43 kDa (*TARDBP*), the gene encoding TDP-43, are a rare cause of ALS, ∼97% of ALS cases and ∼50% of patients with frontotemporal dementia (FTD) present with TDP-43 proteinopathy characterized by TDP-43-positive insoluble nuclear and cytoplasmic inclusions in affected neurons (Arai et al., 2006; Guo and Shorter, 2017; Neumann et al., 2006).

TDP-43 is an essential, highly conserved, ubiquitously expressed, and predominantly nuclear protein with RNA/DNA-binding properties (Portz et al., 2021). It acts as a transcriptional repressor and is implicated in RNA transport and stability, alternative splicing, microRNA biogenesis, and formation of stress granules (SGs) (Alami et al., 2014; Aulas et al., 2012; Buratti and Baralle, 2008; Kawahara and Mieda-Sato, 2012; Khalfallah et al., 2018; Lalmansingh et al., 2011; Li et al., 2018; McDonald et al., 2011; Tollervey et al., 2011). TDP-43 is comprised of an N-terminal domain involved in dimerization and recruitment of other RNA-binding proteins, a bipartite nuclear localization sequence (NLS), two RNA-recognition motifs (RRM1 and RRM2) and a C-terminal glycine-rich, low complexity prion-like domain (PrLD) that mediates protein–protein interactions and formation of membraneless organelles, such as SGs, through liquid–liquid phase separation (LLPS) (Figure 1A) (Ayala et al., 2008; Buratti et al., 2005; Chang et al., 2012; Freibaum et al., 2010; Jiang et al., 2017; Johnson et al., 2009; Li et al., 2018; Molliex et al., 2015; Winton et al., 2008). The majority of ALS-linked TDP-43 mutations reside in the PrLD, which can enhance aggregation propensity and also indicates that this region of TDP-43 may be prone to pathological protein modifications (Figure 1A) (Harrison and Shorter, 2017; Johnson et al., 2009; Prasad et al., 2019).

**Figure 1.**
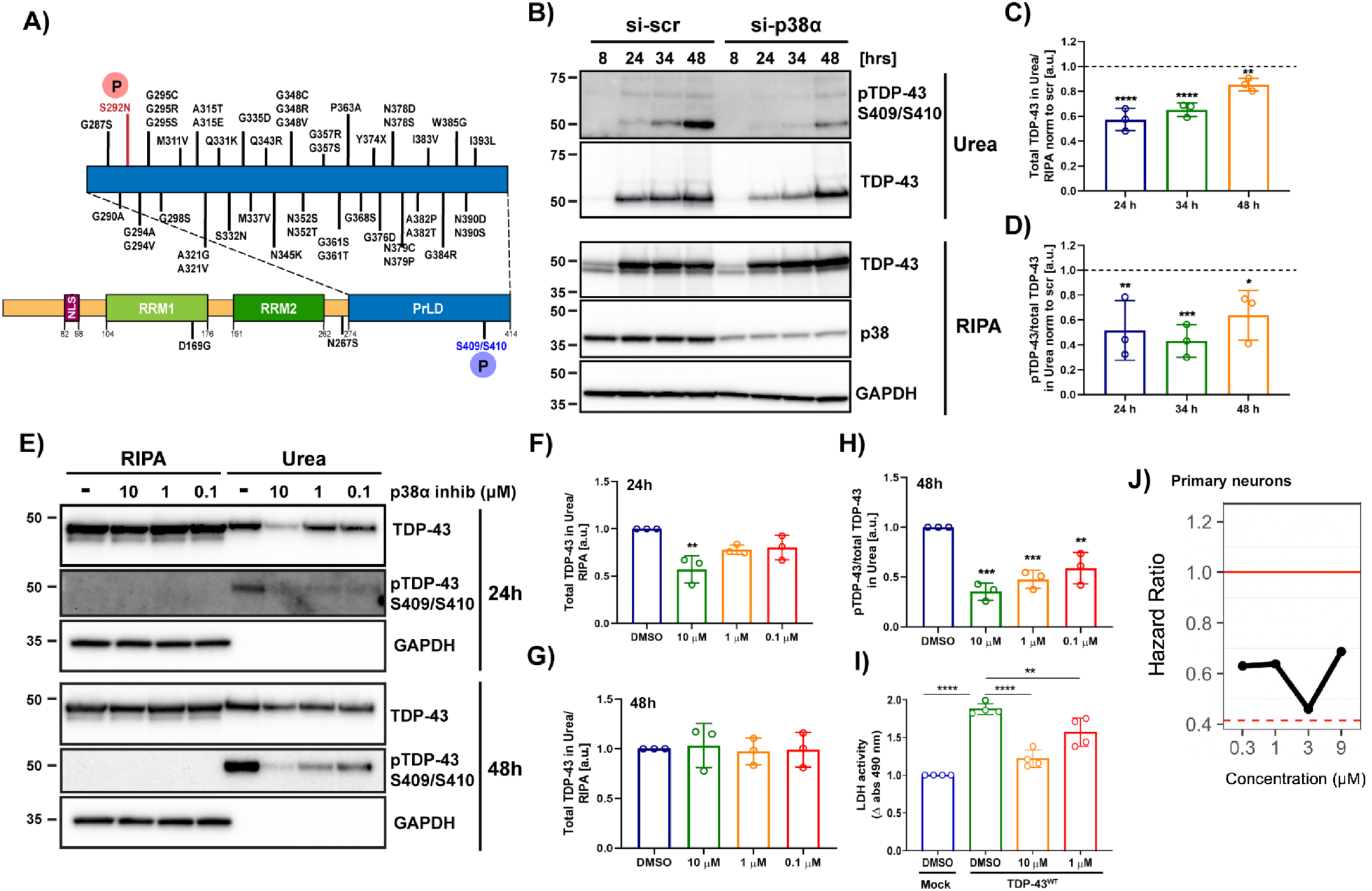
Genetic and pharmacological inhibition of p38α MAPK reduces TDP-43 aggregation and phosphorylation at S409/S410, and inhibits TDP-43-induced neurotoxicity. **A)** Schematic of domain architecture of TDP-43 and location of ALS-linked mutations and phosphorylation sites (P) detected after p38α MAPK treatment *in vitro*. **B)** Western blot of total and phosphorylated TDP-43^M337V^ in RIPA and urea fractions of SH-SY5Y cells with siRNA-induced p38α knockdown. GAPDH was used as a loading control. **C)** Quantification of urea/RIPA ratio of total TDP-43 normalized to levels in scrambled siRNA (si-scr). **D)** Quantification of pTDP-43/total TDP-43 ratio in urea fraction normalized to levels in si-scr (mean band signal ± SD, two-way ANOVA with Sidak’s multiple comparison test, n = 3). **E)** Western blot of total and pTDP-43^M337V^ in RIPA and urea fractions of SH-SY5Y cells with pharmacological p38α inhibition with compound 1. GAPDH was used as a loading control. Quantification of urea/RIPA ratio of total TDP-43 at time points 24h **(F)** and 48h **(G)** post-transfection, and pTDP-43/total TDP-43 ratio in urea fraction at time point 48h post-transfection **(H)** normalized to levels in DMSO-treated cells (mean band signal ± SD, one-way ANOVA with Dunnett’s multiple comparison test, n = 3). **I)** Quantification of LDH activity in conditioned medium normalized to levels in DMSO-treated TDP-43^WT^ -transfected NSC-34 cells. (one-way ANOVA with Dunnett’s multiple comparison test, n = 4 with 6 replicates in each). ^∗^p < 0.05, ^∗∗^p < 0.01 ^∗∗∗^p < 0.001, ^∗∗∗∗^p < 0.0001. **J)** Hazard ratios of primary neurons expressing mApple and TDP-43^M337V^-EGFP treated with p38α inhibitor VX-745 at 0.3, 1, 3, and 9 µM compared with DMSO control (reference, set at 1.0) were 0.6296, 0.6378, 0.4596, and 0.6869 respectively. Reduction of hazard ratio was most significant at 3 µM (Cox proportional hazard, p<0.01). See also Figure S1.

Phosphorylation at serine residues 409/410 (S409/S410) is one of the major pathological markers for TDP-43 inclusions in human brains (Hasegawa et al., 2008; Neumann et al., 2009). In addition to phosphorylation, TDP-43 also undergoes other post-translational modifications (PTMs) in ALS patients and disease-mimicking models, including ubiquitination, generation of C-terminal domain fragments (CTFs), cysteine oxidation, sumoylation, and acetylation (Buratti, 2018). However, the functional and pathological significance of TDP-43 PTMs remains unknown. Thus, a clear understanding of the effects of phosphorylation and other PTMs on solubility, localization, and aggregation propensity of TDP-43 will enable new insights into the mechanisms of ALS pathogenesis.

The mitogen-activated protein kinase (MAPK) signaling pathway plays a key role in the regulation of cellular differentiation, motility, growth, and survival (Brennan et al., 2021). MAPKs, which include extracellular-signal-regulated kinases (ERK), Jun amino-terminal kinases (JNK) and p38 MAPKs, are activated in response to cytokines, growth factors and various stressors including oxidative stress and endoplasmic reticulum stress (Brennan et al., 2021; Cuadrado and Nebreda, 2010; Morrison, 2012). In mammals, p38 MAPKs comprise four isoforms (α, β, γ and δ) (Zarubin and Han, 2005). Of these, α and β are expressed in most tissues, including the brain (Yasuda et al., 2011). p38γ is most highly expressed in skeletal muscle, and p38δ in testis, pancreas, kidney and small intestine (Cuenda et al., 1997). Aberrant p38 signaling has been linked to several neurodegenerative diseases, including ALS (Burton et al., 2021). Specifically, p38 MAPK inhibition can reduce motor neuron apoptosis and restore the physiological rate of axonal retrograde transport in SOD1-ALS models (Dewil et al., 2007; Gibbs et al., 2018; Pickhardt et al., 2019). Furthermore, both genetic and pharmacological inhibition of p38β in a *Drosophila* model of TDP-43 toxicity rescued premature lethality (Zhan et al., 2015).

In this study, we establish a role for p38α MAPK in promoting TDP-43 proteinopathy. We show that inhibition of p38α MAPK reduces ALS-associated TDP-43 phenotypes, including TDP-43 aggregation, S409/S410 phosphorylation, cytoplasmic accumulation, and toxicity. Furthermore, *in vitro* kinase assays combined with mass spectrometry revealed that p38α directly phosphorylates TDP-43 at S292 and S409/S410. Subsequent cellular experiments identified S292, a residue previously found to be altered in a genetic risk allele for ALS (S292N) (Xiong et al., 2010; Zou et al., 2012), as an important site in regulating TDP-43 aggregation. We found that phosphorylation at S292 induced phosphorylation at S409/S410 and promoted TDP-43 aggregation. Moreover, we found that the residue R293, adjacent to S292, is methylated by protein arginine methyltransferase 1 (PRMT1). Interestingly, loss or gain of function for other PRMTs is implicated in the pathogenesis of neurodegenerative diseases (Ratovitski et al., 2015; Simandi et al., 2018). We found that phosphorylation at S292 inhibited R293 methylation, whereas R293 methylation reduced phosphorylation at S409/S410, suggesting an interplay between phosphorylation at S292 and S409/S410 and methylation at R293. Biochemical studies indicated that S292 and S409/S410 phosphorylation reduce TDP-43 LLPS but allow TDP-43 aggregation. By contrast, R293 methylation allows TDP-43 LLPS but reduces TDP-43 aggregation. Taken together, our results reveal additional regulatory mechanisms in TDP-43 homeostasis mediated by p38α and PRMT1. We suggest that strategies aimed at reducing p38α-mediated TDP-43 phosphorylation and promoting R293 methylation could have therapeutic utility for ALS and related TDP-43 proteinopathies.

## Materials and methods

### Plasmids

cDNA sequences were based on the accession number NM_007375.3 for human TARDBP, NM_001315 for human p38*α* (MAPK14) and NM_001536 for human PRMT1. Plasmids harboring N-terminally myc-tagged wild-type or M337V-mutant TDP-43 sequences were generated as described (Wobst et al., 2017). Plasmids harboring C-terminally Flag-tagged wild-type or mutant human TDP-43 and p38*α* sequences in the pcDNA3.1+/C-(K)-DYK mammalian expression vector were purchased from Genscript and PRMT1 plasmid was purchased from Origene. TDP-43 bacterial expression vector harboring a C-terminal MBP tag (pJ4M TDP-43-TEV-MBP-6xHis) was purchased from Addgene (Plasmid # 104480). TDP-43 mutations were generated by site-directed mutagenesis using QuikChange (Agilent) and confirmed by DNA sequencing. The presence of the S409:S410E, S409:S410A, S292E, S292N, S292A, S292:S409:S410A, S292:S409:S410E, and R293F mutations in TDP-43, as well as dominant negative (DN) T180A/Y182F (Winzen et al., 1999) and constitutively active (CA) D176A/F327S mutations (Diskin et al., 2004) in p38*α* were confirmed by DNA sequencing performed by Tufts University Core Facilities (forward sequencing primer 5’-TAA-TAC-GAC-TCA-CTA-TAG-GG-3’, reverse sequencing primer 5’-CAG-GAA-ACA-GCT-ATG-AC-3’) and Eurofins Genomics (TDP-43 sequencing primer 1 5’-TAA-TAC-GAC-TCA-CTA-TAG-GGG-AAT-TG-3’, sequencing primer 2 5’-CGG-TGA-GGT-GCT-GAT-GGT-CC-3’, sequencing primer 3 5’-GGC-TTT-GGC-AAT-TCG-CGT-GG-3’)

### Synthesis of 6-(2,4-difluorophenoxy)-8-ethyl-2-(2-isopropoxyethylamino)pyrido[2,3-d]pyrimidin-7-one (compound 1)

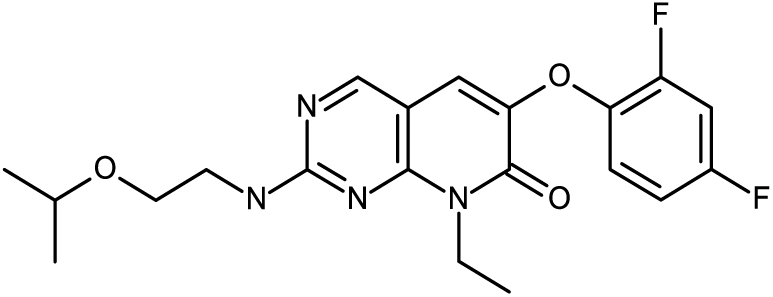

A stirred solution of 2-isopropoxyethan-1-amine (406 mg, 3.93 mmol) was supplemented with 6-(2,4-difluorophenoxy)-8-ethyl-2-(methylsulfonyl)pyrido[2,3-d]pyrimidin-7(8H)-one (300 mg, 0.79 mmol) in DMF (3 mL). The resulting solution was stirred at 80 °C for 1 hour in a microwave reactor. The crude product was purified by preparative HPLC (Column: Xselect CSH OBD Column 30*150mm 5 µm n;Mobile Phase A:Water (0.1%FA), Mobile Phase B: ACN; Flow rate: 60 mL/min; Gradient: 40% B to 72% B in 8 min; 254/220 nm; Rt: 7.57 min). Fractions containing the desired compound were evaporated to dryness to afford 6-(2,4-difluorophenoxy)-8-ethyl-2-((2-isopropoxyethyl)amino)pyrido[2,3-d]pyrimidin-7(8H)-one (190 mg, 59.7 %) as a white solid. ^1^H NMR (400 MHz, DMSO-*d*_6_) δ 8.56 (s, 1H), 7.76 (m, 1H), 7.52 - 7.41 (m, 2H), 7.26-7.18 (m, 1H), 7.06-7.02 (m, 1H), 4.42-4.32 (m, 2H), 3.65-3.45 (m,5H), 1.28-1.22 (m, 3H), 1.09-1.02 (m, 6H). m/z (ES+), [M+H]+ = 405; TFA, HPLC tR = 1.822 min. The intermediate 6-(2,4-difluorophenoxy)-8-ethyl-2-(methylsulfonyl)pyrido[2,3-d]pyrimidin-7(8H)-one was prepared as described (Goldstein et al., 2011). Compound 1 was tested in a Z’-Lyte kinase assay for p38*α* (Thermo Fisher) and inhibited p38*α* activity with an IC_50_ of 25nM.

### Cell culture, transfections and inhibition of p38α MAPK and methyltransferase activity

SH-SY5Y cells were cultured in minimum essential medium (MEM; Thermo Fisher Scientific) supplemented with L-glutamine, 10% fetal bovine serum and 1% penicillin/streptomycin in a humidified incubator at 37°C and 5% CO_2_. For immunocytochemical analysis, cells were grown on glass coverslips coated with 1 mg/mL poly-*L*-lysine (Sigma) in 24-well plates (Cellstar). Cells were transiently transfected using Fugene HD (Promega) following the manufacturer’s instructions 24 hours after seeding (Fugene HD:DNA ratio 3:1). Total transfection times were 8 hours - 48 hours. For pharmacological inhibition of p38α MAPK, compound 1 (synthesized by AstraZeneca) or equal volume of DMSO vehicle, was added to cells for 24 or 48 hours at a final concentration of 0.1 μM, 1 µM or 10 µM. For pharmacological inhibition of arginine methyltransferase activity, adenosine-2′,3′-dialdehyde (AdOx) (Sigma) or equal volume of DMSO vehicle was added to cells for 24 hours at a final concentration of 20 µM.

### Antibodies

The following antibodies were used for immunocytochemical staining: rabbit N-terminal TDP-43 (1:500; 10782-2-AP, Proteintech), mouse anti FLAG-tag (1:500; A00187, GenScript). Anti-rabbit and anti-mouse secondary antibodies coupled to Alexa-488, and Alexa-647 were used for detection (1:1,000; Thermo Fisher Scientific). For western blot analysis, the following antibodies were used: rabbit N-terminal TDP-43 (1:7,000; 10782-2-AP, Proteintech), mouse S409/S410 phospho-TDP-43 (1:3,000; CAC-TIP-PTD-M01, Cosmo), rabbit p38 MAPK (1:1,500; 9212S, Cell Signaling Technology), rabbit COX IV (3E11) (1:1,500; 4850, Cell Signaling Technology), mouse Histone H3 (96C10) (1:1,500; 3638, Cell Signaling Technology), rabbit Mono-Methyl Arginine (R*GG) (D5A12) (1:1,500; 8711S, Cell Signaling Technology), rabbit Asymmetric Di-Methyl Arginine Motif [adme-R] MultiMab (1:1,500; 13522S, Cell Signaling Technology), rabbit Symmetric Di-Methyl Arginine Motif [sdme-RG] MultiMab (1:1,500; 13222S, Cell Signaling Technology), rabbit PRMT1 (A33) (1:1,500; 2449, Cell Signaling Technology), mouse GAPDH (1:10,000; 60004-1-Ig, Proteintech). Anti-mouse and anti-rabbit horseradish peroxidase-coupled secondary antibodies were purchased from Jackson Immunoresearch (1:10,000).

### siRNA knockdown of p38α and PRMT1

Small interfering RNAs (siRNA) were obtained from Thermo Fisher Scientific: p38*α*: s3585 and s3586, PRMT1: s6917, s6919, Negative control: 4390846, 4390843. Reverse transfection was performed with Lipofectamine RNAiMAX-reagent (Invitrogen) following the manufacturer’s instructions. In brief, 25 pmol of siRNA were mixed with 5 µL of Lipofectamine RNAiMAX in 500 µL of Opti-MEM (Invitrogen) in a 6-well plate. The mixture was incubated for 20 min at room temperature and 3-3.5 x 10^5^ cells were added to mixture. Total knockdown times were 8 - 72 hours.

### Sequential extraction of insoluble protein aggregates

Extraction of insoluble proteins was performed as previously described (Wobst et al., 2017). Briefly, transfected cells were lysed in 300 μL radio-immunoprecipitation assay (RIPA) buffer [50 mM Tris-HCl, 150 mM NaCl, 1% NP-40, 0.5% sodium deoxycholate, and 0.1% SDS] (Boston bioproducts) supplemented with 2 mM EDTA, protease inhibitors (cOmplete, Roche), and phosphatase inhibitors (PhosStop, Roche). Lysates were sonicated 2 x 15 s with 20% maximum amplitude and centrifuged for 30 min at 100,000 × *g* and 4°C. The supernatant was collected as the RIPA-soluble fraction. The pellet was washed in RIPA buffer and centrifuged as above. The supernatant was discarded and the urea-soluble fraction was generated by resuspending the pellet in 100 μL urea buffer [7 M urea, 2 M thiourea, 30 mM Tris pH 8.5, 4% 3-[(3-Cholamidopropyl) dimethylammonio]-1-propanesulfonate (CHAPS, Sigma)], and sonicating the samples as above followed by centrifugation at room temperature for 30 min at 100,000 × g. The supernatant was collected as the urea-soluble fraction.

### Immunoblotting

Cell extracts or RIPA and urea fractions obtained from sequential extractions were diluted with NuPAGE sample buffer. Proteins were separated on 4-12% NuPAGE Bis-Tris gels (Thermo Fisher Scientific) under denaturing conditions and transferred onto polyvinylidene difluoride (PVDF) membranes (Millipore). All blocking and antibody incubation steps were performed either in 5% milk in Tris-buffered saline (TBS) [25 mM Tris, 3 mM KCl, 140 mM NaCl, pH 7.4] supplemented with 0.05% Tween-20 (TBS-T) (Thermo Fisher Scientific) or in 5% bovine serum albumin (BSA) in TBS-T. Western blots were developed with enhanced chemiluminescent substrates (ECL). Digital images were acquired with a ChemiDoc MP imaging system (BioRad). Where necessary, blots were stripped with stripping buffer for 15 min (Restore, Thermo Fisher Scientific) and re-probed with loading control antibodies.

### Lactate Dehydrogenase (LDH) assay to monitor cell death

NSC-34 cells were cultured in Dulbecco’s modified Eagle’s medium (DMEM, Thermo Fisher Scientific), supplemented with 10% fetal bovine serum and 1% penicillin/streptomycin in a humidified incubator at 37°C and 5% CO_2_. Cells grown on 96-well plates coated with poly-*L*-lysine (BioCoat multiwell plates, Corning) were transiently transfected using Lipofectamine 3000 (Thermo Fisher Scientific) following the manufacturer’s instructions 24 h after seeding (Lipofectamine 3000:DNA ratio 3:1). Cells were treated with DMSO or p38α MAPK inhibitor (compound 1) at final concentrations of 1 or 10 µM for 4 hours post-transfection. LDH activity was measured from 50 µL of conditioned medium using LDH assay kit (Life Technologies) following the manufacturer’s instructions.

### Longitudinal imaging and neuronal survival analysis

Primary cortical neurons from E17 mouse embryos were cultured in 384-well plates. Neurons were co-transfected with plasmids expressing the fluorescent marker mApple and TDP-43^M337V^-EGFP. Neurons were treated with p38*α* inhibitor, VX-745 (synthesized by AstraZeneca), at 0.3, 1, 3, or 9 µM, and imaged daily for 7 days on a custom-built highthroughput robotic microscopy system (Arrasate and Finkbeiner, 2005; Barmada et al., 2010). Images were montaged and neurons were segemented and tracked using custom-built algorithms. Cox proportional hazard analysis was used to determine hazard ratios. DMSO was used as a control.

### Cytoplasmic and nuclear protein extraction

Cytoplasmic and nuclear protein extraction was performed using a commercial subcellular protein fractionation kit (Thermo Fisher Scientific). Briefly, ∼80% confluent SH-SY5Y cells plated in a 100 -mm dish were transfected as indicated for 24 hours. Cells were trypsinized and rinsed with cold phosphate-buffered saline (PBS), and cell suspensions were transferred to pre-chilled 1.5 mL microcentrifuge tubes. Cells were lysed in cytoplasmic extraction buffer at 4°C for 10 minutes with gentle mixing. After centrifugation at 500 x g at 4 °C for 5 min, the supernatant was collected and stored as the cytoplasmic fraction. After addition of membrane extraction buffer to the pellet, the tube was vortexed vigorously for 5 s and incubated at 4°C for 10 minutes with gentle mixing. After centrifugation at 3,000 x g at 4 °C for 5 min, the supernatant was collected and stored as the membrane fraction. The pellet was resuspended in nuclear extraction buffer followed by vigorous vortexing for 15 s and incubation at 4°C for 30 minutes with gentle mixing. After centrifugation at 5,000 x g at 4 °C for 5 min, the supernatant was collected and stored as the soluble nuclear fraction. The pellet was resuspended in chromatin-bound extraction buffer followed by vigorous vortexing for 15 s and incubation at room temperature for 15 s. The vortexing and incubation steps were repeated twice. After centrifugation at 16,000 x g at 4 °C for 5 min, the supernatant was collected and stored as the chromatin-bound nuclear fraction. Bicinchoninic acid (BCA) assay was used to measure protein concentrations.

### Immunocytochemistry

Cells grown on poly-*L*-lysine-coated glass coverslips were washed in PBS and fixed in 4% paraformaldehyde (in PBS) for 15 min followed by permeabilization in 0.25% Triton-X (in PBS) for 10 min. Cells were blocked with 10% normal goat serum (in PBS, Abcam) for 1 h at room temperature and incubated overnight at 4°C in primary antibody diluted in blocking solution. The next day, cells were washed with PBS and incubated for 1 h in secondary antibody diluted in blocking solution. Coverslips were mounted with Prolong Gold Antifade Mountant with DAPI (Thermo Fisher Scientific). Image acquisition was performed using a Nikon A1 confocal/Eclipse Ti inverted microscope system and NIS Elements software (Nikon).

### Protein kinase assays with recombinant proteins

5 μL of 10x kinase assay buffer [25 mM Tris (pH 7.5), 5 mM β-glycerophosphate, 2 mM DTT, 0.1 mM Na_2_VO_4_, 10 mM MgCl_2_] (Cell Signaling Technology), 200 µM ATP (Cell Signaling Technology), 750 ng of recombinant human TDP-43 protein (Proteintech) and 300 ng of recombinant active p38α kinase (SignalChem) were mixed in pre-chilled 1.5 mL tubes. Reactions were made up to a total volume of 50 μL with ddH_2_O. Samples were mixed by flicking the tubes followed by brief centrifugation at 4°C and incubation at 30°C for 30 min. When indicated, kinase reactions were treated with 10 µM the p38α inhibitor compound 1. Reactions were stopped by adding 18 μL of 4× NuPAGE sample buffer and boiling samples at 95°C for 5 minutes. Samples were subjected to immunoblotting or Coomassie staining followed by phosphorylation analysis by LC-MS/MS.

### Phosphorylation analysis by LC-MS/MS

*In vitro* kinase reactions were separated on 4-12% NuPAGE Bis-Tris gels (Thermo Fisher Scientific) and gel bands were visualized with SimplyBlue™ SafeStain (Thermo Fisher Scientific). Gel bands were excised and cut into ∼1 mm^3^ pieces. The samples were reduced with 1 mM dithiothreitol (DTT) for 30 minutes at 60°C and then alkylated with 5 mM iodoacetamide for 15 minutes at room temperature. Gel pieces were then subjected to a modified in-gel trypsin digestion procedure (Shevchenko et al., 1996). Briefly, gel pieces were washed and dehydrated with acetonitrile for 10 min followed by removal of acetonitrile. Pieces were then completely dried using a speed-vac. Gel pieces were rehydrated in 50 mM ammonium bicarbonate solution containing 12.5 ng/µL modified sequencing-grade trypsin (Promega) at 4°C before incubation overnight at 37°C. Peptides were later extracted by removing the ammonium bicarbonate solution, followed by one wash with a solution containing 50% acetonitrile and 1% formic acid. The extracts were then dried in a speed-vac for 1 hour and stored at 4°C until analysis. On the day of analysis, samples were reconstituted in 5 - 10 µL of High Performance Liquid Chromatography (HPLC) solvent A (2.5% acetonitrile, 0.1% formic acid). A nano-scale reverse-phase HPLC capillary column was created by packing 2.6 µm C18 spherical silica beads into a fused silica capillary (100 µm inner diameter, ∼30 cm length) with a flame-drawn tip (Peng and Gygi, 2001). After equilibrating the column, each sample was loaded onto the column via a Famos auto sampler (LC Packings). A gradient was formed, and peptides were eluted with increasing concentrations of solvent B (97.5% acetonitrile, 0.1% formic acid). As each peptide was eluted it was subjected to electrospray ionization before entering an LTQ Orbitrap Velos Pro ion-trap mass spectrometer (Thermo Fisher Scientific). Eluting peptides were detected, isolated, and fragmented to produce a tandem mass spectrum of specific fragment ions for each peptide. Peptide sequences were determined by matching protein or translated nucleotide databases with the acquired fragmentation pattern using the software program Sequest (Thermo Finnigan) (Eng et al., 1994). The modification of 79.9663 mass units to serine, threonine, and tyrosine was included in the database searches to determine phosphopeptides. Phosphorylation assignments were determined using the Ascore algorithm (Beausoleil et al., 2006). All databases include a reversed version of all sequences and the data was filtered to 1-2% peptide false discovery rate.

### Co-immunoprecipitation (Co-IP)

Approximately 80% confluent SH-SY5Y cells plated in 6-well plates or 100-mm dishes were transfected as indicated. Cells were rinsed with cold PBS, and then lysed in cold lysis buffer [20 mM Tris-HCl (pH 7.5), 150 mM NaCl, 1 mM Na_2_EDTA, 1 mM EGTA, 1% Triton, 2.5 mM sodium pyrophosphate, 1 mM beta-glycerophosphate, 1 mM Na_3_VO_4_, 1 µg/mL leupeptin] (Cell Signaling Technology), supplemented with protease inhibitors (cOmplete, Roche). Cells were incubated on ice for 5 minutes, collected in pre-chilled 1.5 mL tubes, sonicated briefly and cleared by centrifugation (14,000 x g at 4°C for 10 min). Lysate aliquots were stored as the input samples. Cell lysates were either incubated with Anti-FLAG® M2 Magnetic Beads (Millipore Sigma) overnight with continuous rotation at 4°C, or were subjected to pre-cleaning with protein A Dynabeads (Thermo Fisher Scientific) for 1 hour at 4°C followed by incubation with indicated primary antibodies overnight with continuous rotation at 4°C. Protein A Dynabeads were then added to pre-cleared antibody-containing samples, and the incubation was continued for an additional 2 hours at room temperature. Beads with immunoprecipitated proteins were washed 5x with either TBS [50 mM Tris HCl, 150 mM NaCl, pH 7.4] (Anti-FLAG® M2 Magnetic Beads) or lysis buffer (Protein A Dynabeads). Immunoprecipitated proteins were eluted with 2x NuPAGE sample buffer by boiling for 3 min. Both input samples and immunoprecipitated proteins were analyzed by immunoblotting.

### Purification of recombinant TDP-43-MBP

Wild-type TDP-43-MBP-6xHis and TDP-43 mutants S292E, S409:S410E, S292:S409:S410E, and R293F were expressed and purified as previously described (Wang et al., 2018). Briefly, TDP-43 variants were expressed in *E. coli* BL21-CodonPlus (DE3)-RIL cells (Agilent). Cell cultures were grown to an OD_600_ of ∼0.5-0.7 and then cooled down to 16°C. Protein expression was induced with 1 mM IPTG overnight. Cells were harvested and resuspended in purification buffer (20 mM Tris-HCl, pH 8.0, 1 M NaCl, 10 mM imidazole, 10% (v/v) glycerol, 2 mM β-mercaptoethanol supplemented with cOmplete EDTA-free protease inhibitor cocktail) and lysed using 1 mg/mL lysozyme and sonication. Proteins were purified using Ni-NTA agarose (Qiagen) and eluted using 300 mM imidazole in purification buffer. Proteins were then further purified over amylose resin (NEB) and eluted with elution buffer (20 mM Tris-HCl, pH 8.0, 1 M NaCl, 10 mM imidazole, 10 mM maltose, 10% (v/v) glycerol, and 1 mM DTT). Purified proteins were concentrated, flash frozen and stored at -80°C.

### *In vitro* TDP-43 aggregation assay

Purified recombinant TDP-43-MBP-6xHis wild-type and TDP-43 mutants were first thawed and buffer exchanged into 20 mM HEPES-NaOH, pH 7.4, 150 mM NaCl and 1mM DTT using Micro Bio-Spin^TM^ P-6 Gel Columns (Bio-Rad). Protein concentration was determined by NanoDrop, and the final concentration of TDP-43 was then adjusted to 5 µM in the same buffer. To measure aggregation kinetics, aggregation was initiated by cleavage of the MBP-6xHis tag using 1 µg/mL TEV protease (Cupo and Shorter, 2020) at t = 0, and turbidity was measured over 16 h at an absorbance of 395 nm using a TECAN M1000 plate reader. Values were normalized to wild-type TDP-43 + TEV protease to determine the extent of aggregation of TDP-43 mutants.

### *In vitro* TDP-43 LLPS assay

Purified recombinant TDP-43-MBP-6xHis wild-type and TDP-43 mutants were thawed and buffer exchanged as described above for the aggregation assay. Protein concentration was determined by NanoDrop, and reactions were prepared in phase separation buffer (20 mM HEPES-NaOH, pH 7.4, 150 mM NaCl, 1mM DTT, 100 mg/mL dextran from *Leuconostoc spp.* (Sigma)). Protein was always added last to each phase separation reaction, at a final concentration of 10 µM. Reactions were incubated for 30 min at room temperature, and then 7.5 µL of each reaction was mounted onto a glass slide and imaged by differential interference contrast (DIC) microscopy.

### Droplet image analysis

DIC images of TDP-43 wild-type and its variants were analyzed using custom-written code in MATLAB. Each image was first converted into grayscale for processing. Then, Roberts gradient was used to filter out the noise. The pixel weight for each pixel in the image was determined based on the grayscale intensity differences before segmenting the images. The threshold for image segmentation was adjusted manually to ensure complete and accurate conversion to logical array. Droplets in the images were identified using circle Hough transform. The sensitivity was toggled either to detect missed droplets or to reduce the number of false positives. The detected circles were visualized on the original images. Various quantitative parameters, including the average area, total area, number of droplets and the lists of areas, were given as outputs to the code and were analyzed further through GraphPad Prism. The accuracy of the circule Hough transform was limited for droplets that were smaller than 5 pixels, where 10.8 pixels are equivalent to 1µm.

### Quantification of immunocytochemistry and western blots, and statistical analysis

Western blot band densities and immunocytochemical staining were quantified with ImageJ-Win64 software. For statistical analysis, we used GraphPad Prism 7 and 8, and used unpaired t-test or one-way ANOVA followed by Sidak’s or Dunnett’s multiple comparison test, as indicated for each experiment. All assays were repeated at least three times. A *p*-value less than 0.05 was considered statistically significant.

## Results

### Inhibition of p38α reduces TDP-43 aggregation, phosphorylation, and toxicity

Several lines of evidence point to a role for p38 MAPK in the development and progression of ALS (Bendotti et al., 2004; Corrêa and Eales, 2012; Gibbs et al., 2018; Tortarolo et al., 2003; Zhan et al., 2015). Thus, we first asked whether p38α modulates the formation of insoluble TDP-43 aggregates in human neuronal SH-SY5Y cells. Increased expression of wild-type or ALS-linked TDP-43 variants elicits ALS-like phenotypes in various *in vitro* and *in vivo* models (Arnold et al., 2013; Johnson et al., 2009; Watkins et al., 2020; Wobst et al., 2017; Xu et al., 2011). Indeed, elevated expression of wild-type TDP-43 is connected with FTD (Gitcho et al., 2009) and disease-linked TDP-43 aggregation is proposed to increase TDP-43 expression due to loss of TDP-43 autoregulation (Gasset-Rosa et al., 2019; Polymenidou et al., 2011). TDP-43^M337V^, a pathological mutant form of TDP-43 associated with fALS, has been shown to be especially aggregation-prone and highly phosphorylated (Johnson et al., 2009; Sreedharan et al., 2008; Wobst et al., 2017). We therefore used the TDP-43^M337V^ variant in our experiments to detect more prominent changes in TDP-43 solubility and phosphorylation, and to maximize our experimental signal window. Using siRNA-mediated knockdown of p38α and TDP-43^M337V^ expression, we monitored the accumulation of total and phosphorylated TDP-43 (pTDP-43) in RIPA- or urea-soluble fractions over time. We found that the depletion of p38α significantly reduced the accumulation of insoluble TDP-43^M337V^ in the urea fraction at all time points (Figure 1B-1C). Thus, reduction of p38α reduces TDP-43 aggregation in human neuronal cells.

Phosphorylation of S409 and S410 of TDP-43 is a consistent feature in all sporadic and familial forms of TDP-43 proteinopathies (Neumann et al., 2009). TDP-43 is phosphorylated at S409/S410 in pathological inclusions, but S409/S410 phosphorylation is not observed under physiological conditions in the nucleus (Neumann et al., 2009). Interestingly, we observed a marked decrease in pTDP-43 (i.e. TDP-43 phosphorylated at S409/S410) in the urea fraction when p38α is knocked down (Figure 1B and 1D). Similarly, increased expression of TDP-43^WT^, in place of TDP-43^M337V^, or using a different siRNA to knockdown p38α also decreased pTDP-43 in the urea fraction (Figure S1). Thus, reduction of p38α reduces TDP-43 aggregation and pathological phosphorylation of S409/S410 in human neuronal cells.

Next, we assessed whether pharmacological inhibition of p38α activity affects the solubility and the phosphorylation status of TDP-43 in a similar manner to genetic ablation. Indeed, treatment of SH-SY5Y cells with the p38α inhibitor, compound 1 (see Materials and Methods), significantly reduced TDP-43^M337V^ phosphorylation and aggregation, in a concentration- and time-dependent manner (Figure 1E-1H). Note that we only observe TDP-43 phosphorylation at S409/S410 in the insoluble urea fraction and not in the soluble RIPA fraction (Figure 1E), indicating the pathological nature of S409/S410 phosphorylation. Our findings suggest that p38α inhibition is an effective strategy to reduce pathological TDP-43 aggregation and phosphorylation.

To test whether p38α pharmacological inhibition affects toxicity induced by increased TDP-43 expression, we performed an LDH cytotoxicity assay in mouse motor-neuron-like NSC-34 cells. Expression of TDP-43^WT^ (Figure 1I) or TDP-43^M337V^ in NSC-34 cells induces cytotoxicity. Importantly, inhibition of p38α by compound 1 significantly rescued TDP-43^WT^-induced cytotoxicity (Figure 1I). Thus, pharmacological inhibition of p38α mitigates TDP-43 toxicity in motor-neuron-like NSC-34 cells.

To determine whether p38α pharmacological inhibition rescues TDP-43-induced neurodegeneration, we employed a longitudinal imaging system to monitor neuronal survival (Barmada et al., 2010). We cultured primary mouse cortical neurons, uniformly labeled them with a fluorescent protein (mApple), and expressed TDP43^M337V^. We imaged and tracked the neurons daily for 7 days and used Cox proportional hazard analysis to measure the cumulative risk of death and hazard ratios. We have previously shown that this assay is very sensitive to detecting TDP-43-induced neurodegeneration (Barmada et al., 2010). We found that treatment of neurons with a brain-penetrant p38α pharmacological inhibitor, VX-745 (also known as Neflamapimod) (Duffy et al., 2011), which is in phase 2 clinical trials for Alzheimer’s disease, Huntington’s disease, and dementia with Lewy bodies (Germann and Alam, 2020; Prins et al., 2021), significantly reduced the hazard ratio of TDP-43^M337V^-expressing neurons (Figure 1J). Thus, clinical stage, brain-penetrant p38α inhibitors can also mitigate TDP-43 neurotoxicity. Taken together, our results demonstrate that p38α inhibition reduces the pathological aggregation and phosphorylation of TDP-43, and mitigates TDP-43 toxicity in multiple settings.

### Constitutively active p38α promotes TDP-43 aggregation, S409/S410 phosphorylation, and cytoplasmic accumulation

The siRNA knockdown experiments demonstrated that p38α depletion reduces TDP-43 aggregation and phosphorylation at S409/S410. Therefore, we hypothesized that p38α overexpression would increase TDP-43 aggregation and phosphorylation. To test our hypothesis, we co-expressed TDP-43^M337V^ with wild-type, constitutively active (CA), or dominant negative (DN) forms of p38α in SH-SY5Y cells. Expression of WT or DN-p38α did not have any significant effect on S409/S410 phosphorylation or accumulation of TDP-43 in the urea fraction (Figure 2A-2D). By contrast, CA-p38α promoted TDP-43^M337V^ aggregation, as evidenced by the accumulation of TDP-43 in the urea fraction (Figure 2A-2D). CA-p38α also significantly increased TDP-43 phosphorylation at S409/S410 (Figure 2A-2D). Thus, p38α activation promotes pathological TDP-43 phosphorylation and aggregation.

**Figure 2.**
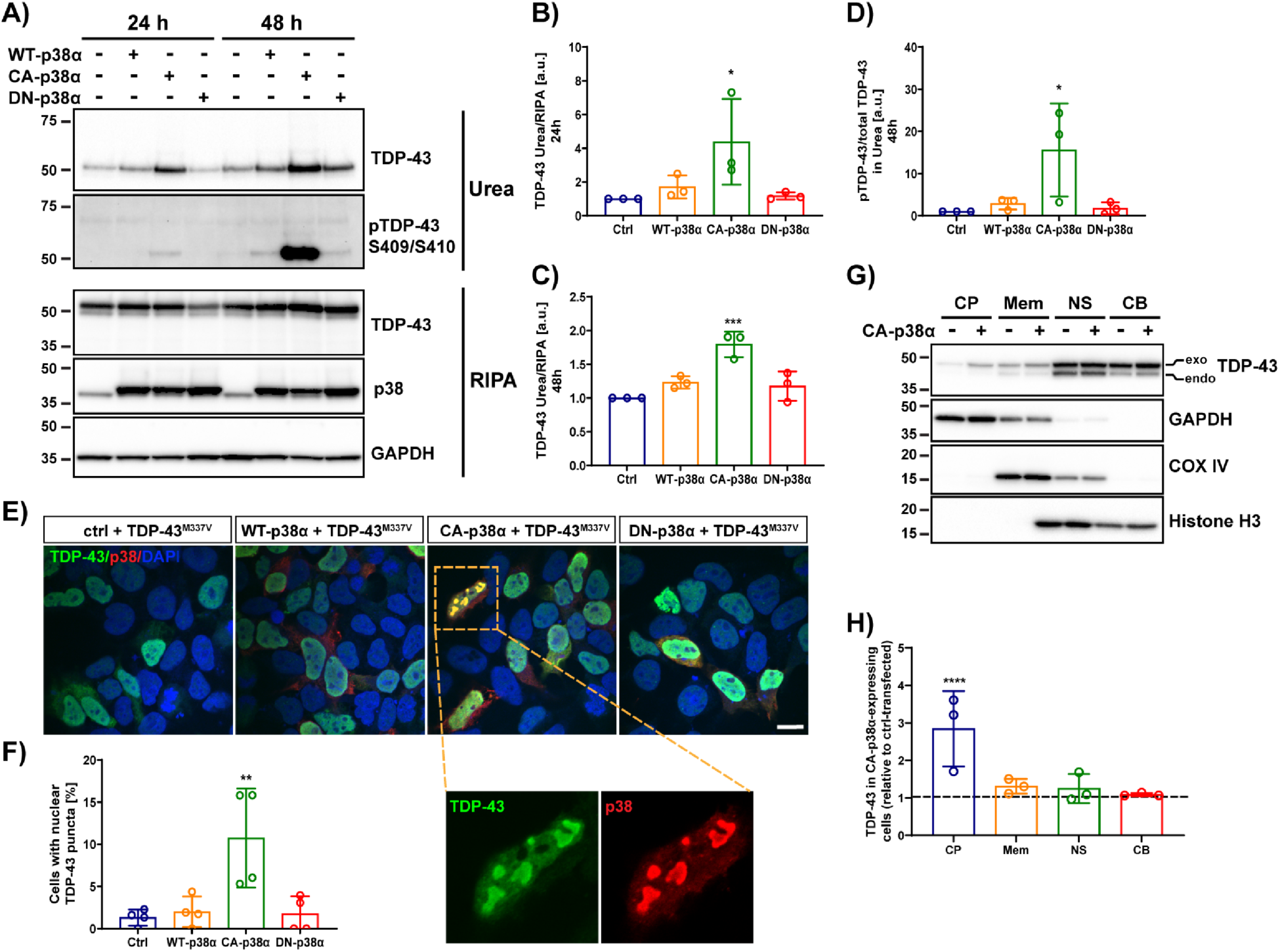
Constitutively active p38α induces aggregation, phosphorylation and cytoplasmic accumulation of TDP-43. **A)** Western blot of total and pTDP-43 in RIPA and urea fractions of SH-SY5Y cells co-transfected with TDP-43^M337V^ and empty control plasmid (Ctrl), WT-p38α, CA-p38α or DN-p38α. GAPDH was used as a loading control. Quantification of urea/RIPA ratio of total TDP-43 at time points 24h **(B)** or 48h **(C)** post-transfection, and pTDP-43/total TDP-43 ratio in urea fraction at time point 48h post-transfection **(D)**, normalized to levels in ctrl-plasmid transfected cells (mean band signal ± SD, one-way ANOVA with Dunnett’s multiple comparison test, n = 3). **E)** Representative confocal images of SH-SY5Y cells co-expressing TDP-43^M337V^ (green) and WT-p38α, CA-p38α or DN-p38α (red) with quantification of the number of cells with nuclear TDP-43 puncta/granules **(F)** (one-way ANOVA with Dunnett’s multiple comparison test, n = 4 and 10 fields each, scale bar 20 µm). **G)** Western blot of TDP-43 in different cellular compartments of SH-SY5Y cells co-transfected with TDP-43^M337V^ and ctrl-plasmid or CA-p38α. Exo denotes exogenous TDP-43 and endo denotes endogenous TDP-43. GAPDH was used as a loading control for cytoplasmic fraction (CP), COX IV for the membrane fraction (Mem), and histone H3 for the soluble nuclear (NS) and chromatin-bound (CB) fractions. **H)** Quantification of total TDP-43 in cytoplasmic, membrane, soluble nuclear and chromatin-bound fractions (mean band signal ± SD, two-way ANOVA with Sidak’s multiple comparison test, n = 3). ^∗^p < 0.05, ^∗∗^p < 0.01 ^∗∗∗^p < 0.001, ^∗∗∗∗^p < 0.0001.

Next, we assessed whether TDP-43^M337V^ co-localizes with p38α in cells. Previously, TDP-43^M337V^ and p38α have been detected in ubiquitinated inclusions (Bendotti et al., 2004; Wobst et al., 2017). Interestingly, we found a significant increase in the number of cells displaying intranuclear TDP-43^M337V^ inclusions specifically when CA-p38α was co-expressed (Figure 2E-2F). Furthermore, these intranuclear TDP-43 inclusions stained positive for p38α (Figure 2E). We also found that expression of CA-p38α promoted the cytoplasmic accumulation of TDP-43, as shown by an increase of TDP-43 in the cytoplasmic fraction (Figure 2G-2H). Together, our data demonstrate that aberrant activation of p38α promotes several hallmarks of ALS pathology, including TDP-43 aggregation, phosphorylation at S409/S410, and cytoplasmic accumulation.

### TDP-43 is directly phosphorylated by p38α at residues S292, S409, and S410

Co-localization of TDP-43 and CA-p38α in intranuclear aggregates might indicate a physical interaction between TDP-43 and p38α (Figure 2E). To explore this possibility, we transfected SH-SY5Y with TDP-43^WT^ and p38α, and then immunoprecipiated p38α or TDP-43 (Figure 3A). TDP-43^WT^ and p38α co-immunoprecipitated in both reciprocal immunoprecipitation experiments, suggesting a robust interaction between TDP-43 and p38α (Figure 3A). Consistently, siRNA-mediated knockdown of p38α reduced the amount of pulled down pTDP-43, and also diminished the amount of endogenous p38α that co-immunoprecipitated with TDP-43^M337V^ (Figure S2), further validating an interaction between p38α and TDP-43.

**Figure 3.**
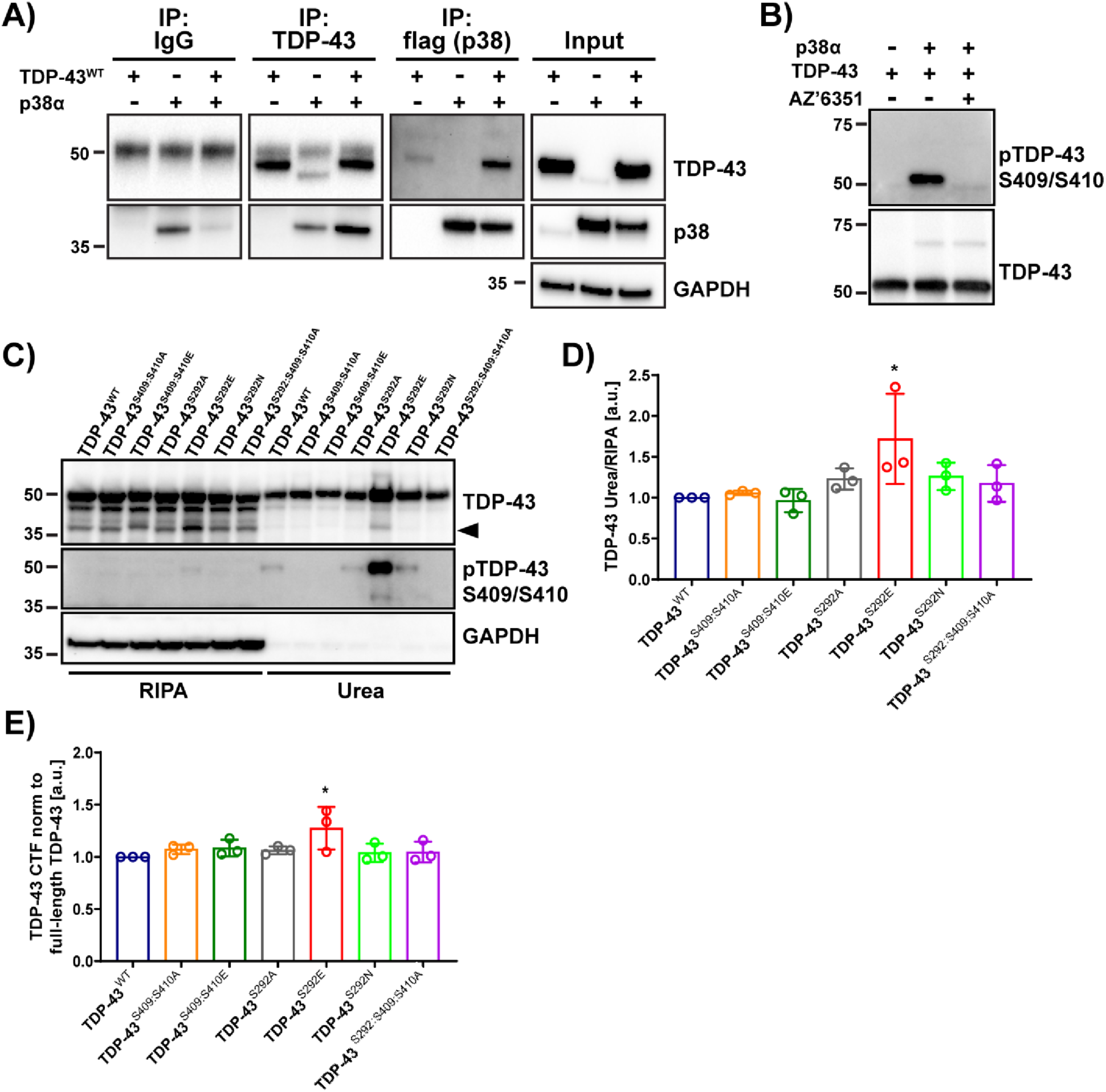
TDP-43 directly interacts with and is phosphorylated by p38α at S292 and S409/S410, with S292 regulating phosphorylation at S409/S410. **A)** Western blot of immunoprecipitated TDP-43^WT^ and co-immunoprecipitated WT-p38α and immunoprecipitated WT-p38α and co-immunoprecipitated TDP-43^WT^ in SH-SY5Y cells. Inputs are shown on the right. GAPDH was used as a loading control. **B)** Western blot of *in vitro* kinase assay (30°C for 30 min) of recombinant human TDP-43 (750 ng) and recombinant active p38α (300 ng) with and without the p38α inhibitor compound 1. **C)** Western blot of total and pTDP-43 in RIPA and urea fractions of SH-SY5Y cells transfected with TDP-43^WT^ and with S292 and S409/S410 mutant constructs. Quantification of urea/RIPA ratio of total TDP-43 **(D)**, and CTF/full length ratio of TDP-43 in RIPA fraction **(E)** at time point 24h post-transfection normalized to levels in TDP-43^WT^ (mean band signal ± SD, one-way ANOVA with Dunnett’s multiple comparison test, n = 3). ^∗^p < 0.05. See also Figure S2.

Next, we asked whether TDP-43 is directly phosphorylated by p38α. Thus, we performed *in vitro* kinase assays. Samples containing recombinant wild-type TDP-43 with or without active p38α were analyzed by western blot analysis after a 30-minute incubation period. In the absence of p38α, no phosphorylation was detected at the S409/S410 site (Figure 3B). However, in the presence of active kinase, a robust band of phosphorylated TDP-43 was observed, indicating that TDP-43 is directly phosphorylated by p38α at S409/S410 (Figure 3B). As expected, the phosphorylation of TDP-43 was prevented by pharmacological inhibition of p38α with compound 1 (Figure 3B).

While S409/S410 residues are thought to be the major pathological phosphorylation sites in TDP-43 (Neumann et al., 2009), we sought to investigate whether p38α phosphorylates TDP-43 at additional serine or threonine residues. Therefore, we performed LC-MS/MS analysis of trypsin-digested TDP-43 after incubation with p38α, which revealed that p38α also phosphorylates TDP-43 at serine residue 292 (S292) (Figure 1A). In fact, TDP-43 is phosphorylated at S292 in the brains of ALS patients (Kametani et al., 2016). Furthermore, a serine-to-asparagine mutation at this site (S292N) is genetically linked to sALS and fALS (Harrison and Shorter, 2017; Xiong et al., 2010; Zou et al., 2012). To investigate the potential effects of phosphorylation at S292 on the aggregation propensity of TDP-43, we took a site-directed mutagenesis approach to generate phospho-mimetic (S292E; TDP-43^S292E^), phospho-dead (S292A; TDP-43^S292A^), and ALS-linked (S292N; TDP-43^S292N^) TDP-43 mutants, which were transfected into SH-SY5Y cells. We compared the aggregation propensity of these S292 mutants to that of TDP-43^WT^ and TDP-43 variants harboring S409:S410A (TDP-43^S409:S410A^), S409:S410E (TDP-43^S409:S410E^) and S292:S409:S410A (TDP-43^S292:S409:S410A^) mutations. Interestingly, overexpressing the phospho-mimetic mutant TDP-43^S292E^ increased the aggregation propensity of TDP-43, as evidenced by the accumulation of TDP-43 in the urea fraction, and also enhanced its phosphorylation at serine residues 409/410 (Figure 3C and 3D). Moreover, TDP-43^S292E^ enhanced the formation of 35 kDa C-terminal fragments of TDP-43 (CTF), which are associated with TDP-43 proteinopathy (Buratti, 2018) (Figure 3C and 3E). Neither TDP-43^S292A^, TDP-43^S292N^, TDP-43^S409:S410A^, TDP-43^S409:S410E^ nor TDP-43^S292:S409:S410A^ had any significant effect on TDP-43 aggregation propensity compared to TDP-43^WT^ (Figure 3C-3D). These results suggest that aggregation of TDP-43 may be enhanced by S292 phosphorylation in neuronal cells. Moreover, S292 phosphorylation also likely stimulates phosphorylation at S409/410.

### TDP-43 undergoes arginine methylation catalyzed by PRMT1

Our data suggest a role for S292 in regulating TDP-43 aggregation via p38α-mediated phosphorylation. An alignment of TDP-43 amino acid sequences revealed that S292 is highly conserved across the phylogenic spectrum from *Homo sapiens* to *Gallus gallus* (Figure 4A). Intriguingly, an arginine-glycine-glycine (RGG) motif directly follows the S292 residue and is also highly conserved (Figure 4A). RGG/RG motifs are preferred substrates for methylation by members of the PRMT family (Thandapani et al., 2013). Given that several studies have shown that arginine methylation can attenuate phosphorylation at nearby residues on the same protein (Guo et al., 2010; Hsu et al., 2011; Lu et al., 2017; Yamagata et al., 2008), we hypothesized that arginine methylation at R293 could potentially interfere with p38α-mediated phosphorylation at the adjacent residue S292. Recent proteomic studies have found that TDP-43 is methylated in HCT116 and HEK293 cells as well as in mouse embryos and human brain tissue (Guo et al., 2014; Larsen et al., 2016). To assess TDP-43 arginine methylation in SH-SY5Y cells, we immunoprecipitated endogenous TDP-43 and probed using antibodies against monomethyl-arginine (MMA), asymmetric dimethyl-arginine (ADMA) or symmetric dimethyl-arginine (SDMA). Western blot analysis showed that immunoprecipitated TDP-43 was indeed mono- and asymmetrically dimethylated and was efficiently recognized by the MMA and ADMA antibodies, but not a SDMA antibody (Figure 4B). Critically, when we depleted PRMT1, the most abundant PRMT in mammalian cells (Tang et al., 2000), using siRNA-mediated knockdown in SH-SY5Y cells, followed by TDP-43 immunoprecipitation and western blot analysis, we found a significant decrease in methylation of endogenous TDP-43 (Figure 4C). Additionally, treating cells with AdOx, a global methyltransferase inhibitor, for 24 hours significantly decreased the methylation of overexpressed TDP-43^WT^, evident by a decrease in immunoprecipitated TDP-43 detected by anti-MMA and anti-ADMA (Figure 4D). Collectively, these data suggest that PRMT1 methylates TDP-43.

**Figure 4.**
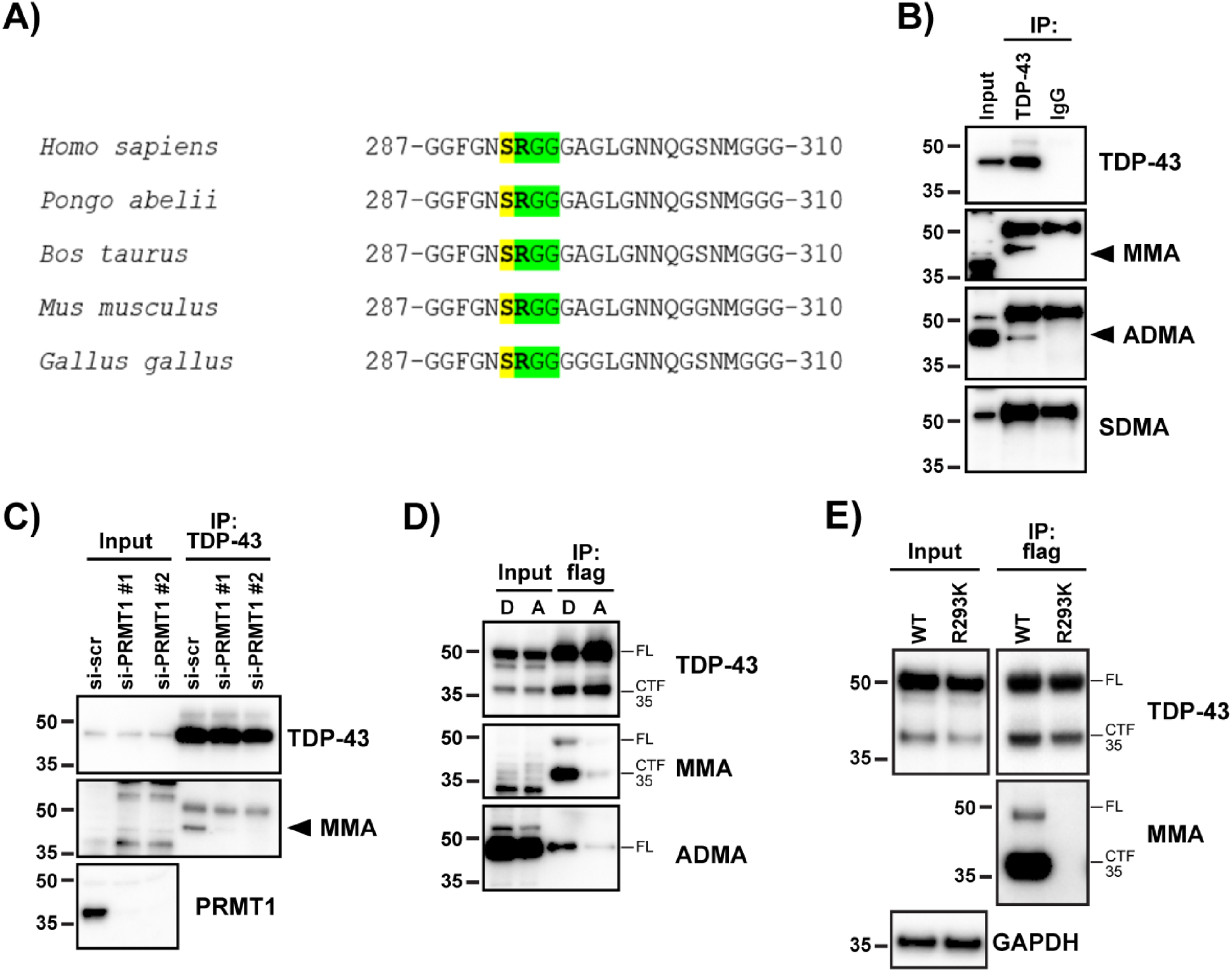
TDP-43 is methylated by PRMT1 primarily at residue R293. **A)** Sequence alignment of amino acids 287-310 of TDP-43 from diverse vertebrate species. Conserved S292 and R293 sites are bolded. S292 phosphorylation site and R293-G295 RGG-motif are highlighted in yellow and green, respectively. **B)** Western blot of immunoprecipitated endogenous TDP-43 from SH-SY5Y-cells probed with antibodies against TDP-43, ADMA, MMA and SDMA. TDP-43 with arginine methylation is marked by arrows. **C)** Western blot of immunoprecipitated endogenous TDP-43 from SH-SY5Y-cells with siRNA-induced PRMT1 knockdown probed with antibodies against TDP-43, MMA and PRMT1. **D)** Western blot of expressed and immunoprecipitated TDP-43^WT^ using anti-Flag antibody from SH-SY5Y-cells treated with DMSO (D) or with methyltransferase inhibitor AdOx (A) at a final concentration of 20 µM for 24h. FL denotes full-length TDP-43 and CTF35 denotes C-terminal 35kDa fragment of TDP-43. **E)** Western blot of expressed and immunoprecipitated TDP-43^WT^ and TDP-43^R293K^ using anti-Flag antibody from SH-SY5Y cells probed with antibodies against TDP-43 and MMA.

Next, we wanted to verify whether R293 is the targeted residue for arginine methylation. Using site-directed mutagenesis, we generated a methylation-dead mutant of TDP-43 by substituting the arginine residue at 293 to a lysine (TDP-43^R293K^). Unlike for TDP-43^WT^, immunoprecipitating TDP-43^R293K^ and probing for monomethylated TDP-43 using anti-MMA antibody revealed an absence of monomethylated TDP-43 (Figure 4E). Overall, our results provide evidence that PRMT1 methylates TDP-43 at R293 in human neuronal cells.

### TDP-43 arginine methylation favors normal LLPS over aberrant aggregation

Arginine methylation of TDP-43 has been largely unexplored. Thus, little is known about R293 methylation and whether it affects TDP-43 LLPS or aggregation propensity at the pure protein level. Likewise, we have limited understanding of how specific phosphorylation events might affect TDP-43 LLPS and aggregation at the pure protein level. To address how phosphorylation and methylation can alter TDP-43 LLPS behavior, we first performed *in vitro* droplet formation assays. Purified recombinant maltose-binding protein (MBP)-tagged TDP-43^WT^, the phospho-mimicking mutants TDP-43^S292E^, TDP-43^S409:S410E^, TDP-43^S292:S409:S410E^, and the arginine methylation-mimic TDP-43^R293F^ (Campbell et al., 2012; Huq et al., 2006; Liu et al., 2019), were separately incubated at physiological concentration (10µM) (Ling et al., 2010) in phase separation buffer containing physiological salt concentration and 10% (w/v) dextran to mimic the crowded cellular environment (Mann et al., 2019; McGurk et al., 2018). Formation of TDP-43 droplets was then visualized using DIC microscopy. TDP-43^WT^ formed spherical droplets that were relatively large in size (average area of ∼12.6µm^2^ ± 1.3), and capable of fusion events indicating liquid-like properties (Figure 5A-C). In contrast, all three phosphomimetic TDP-43 mutants partitioned into droplets that were smaller in size (TDP-43^S292E^, ∼5.8µm^2^ ± 0.6; TDP-43^S409:S410E^, ∼3.9µm^2^ ± 0.3; and TDP-43^S292:S409:S410E^, ∼3.2 µm^2^ ± 0.2) compared to those of TDP-43^WT^ (Figure 5A-C). There was no change in the number of droplets between mutants and TDP-43^WT^. Thus, these reductions in droplet sizes suggest that phosphorylation of S292, S409, and S410 may limit the LLPS propensity of TDP-43. Interestingly, the arginine-methylation mimic, TDP-43^R293F^, formed large droplets that were comparable to TDP-43^WT^ (∼12.6µm^2^ ± 1.9) (Figure 5A-C). These results indicated that phosphorylation and methylation likely have contrasting effects on LLPS of TDP-43. Thus, R293 methylation likely permits wild-type levels of TDP-43 LLPS, whereas S292, S409, and S410 phosphorylation likely reduce TDP-43 LLPS.

**Figure 5.**
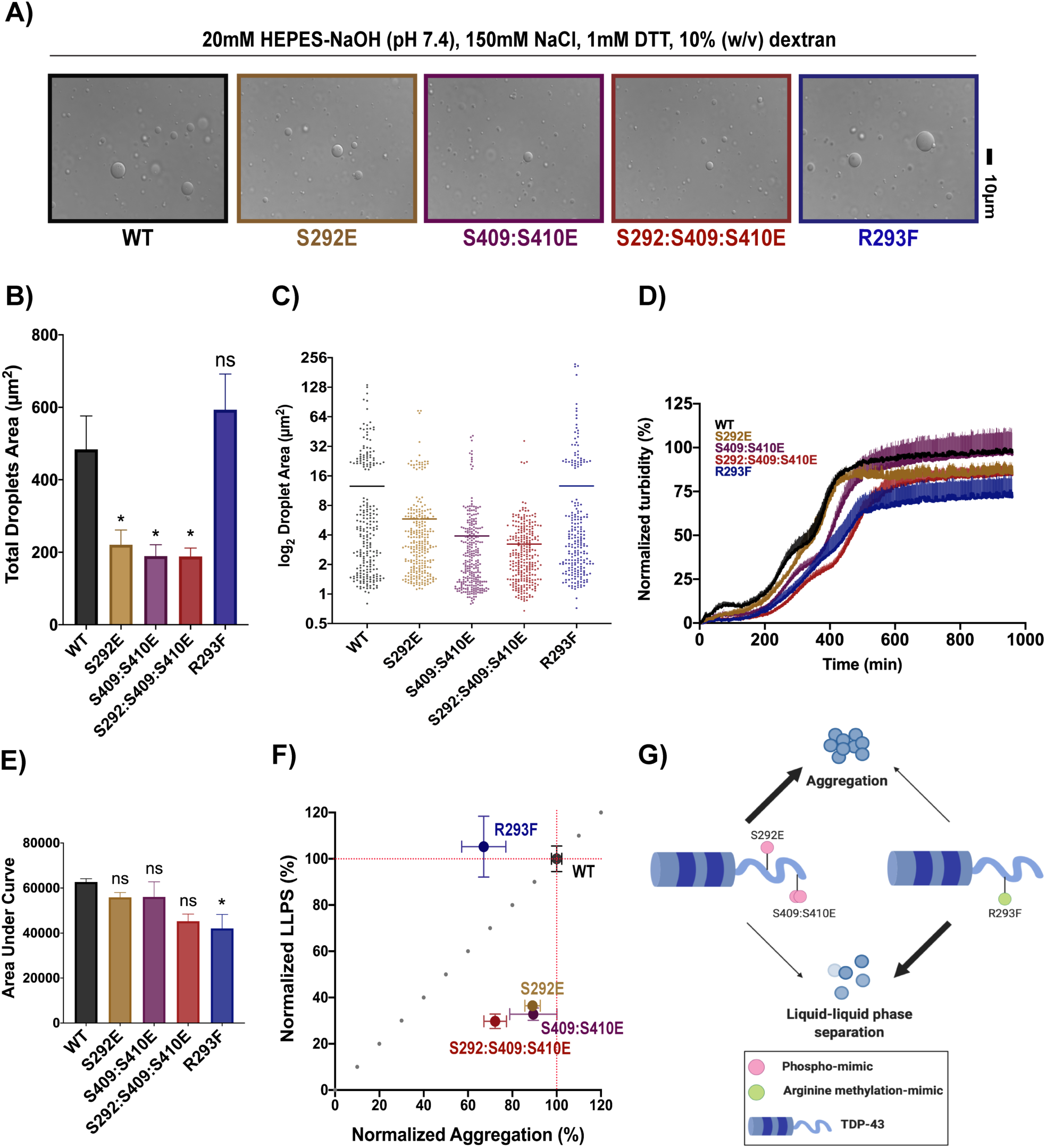
TDP-43 phosphorylation favors aggregation whereas arginine methylation favors LLPS *in vitro.* **A)** Representative DIC microscopy images of liquid-like droplets of 10µM TDP-43-MBP wild-type protein and its mutant forms. Purified recombinant proteins were incubated for 30 minutes with phase separation buffer prior to imaging. Scale bar = 10 µm (n=3). **B)** Bar graph showing the total droplets area for each protein. Mean± SEM, one-way ANOVA with Dunnett’s multiple comparison test (n=3, *p<0.05). **C)** Vertical scatterplot displaying the size distribution of droplets for each protein. Each data point corresponds to a single droplet. Bolded bars represent the average droplet area for each variant. **D)** Turbidity measurements of 5 µM TDP-43-MBP co-incubated with TEV protease (1 µg/mL). Turbidity was measured at an absorbance of 395 nm. Values represent the normalized mean ± SEM (n = 4). **E)** Aggregation data from **D** was quantified by calculating the area under the curve.Values represent means ± SEM (n = 4). One-way ANOVA with Tukey’s multiple comparison test was performed (*p<0.05). **F)** LLPS versus aggregation plot shows the phospho-mimetics cluster below the y=x dotted line, whereas the arginine methylation-mimic appears above the y=x dotted line. Phospho-mimetics inhibit TDP-43 LLPS more severely than TDP-43 aggregation, whereas the arginine-methylation mimic inhibits TDP-43 aggregation but not TDP-43 LLPS. Y-axis represents normalized LLPS propensity relative to wild-type based on the average area of droplets from analysis in **C**. X-axis represents normalized aggregation relative to wild-type from **D**. Error bars represent SEM with n=3 for their respective dimension. **G)** Schematic diagram describing the dichotomy in outcomes between p38*α*-mediated TDP-43 phosphorylation and PRMT1-mediated arginine methylation. Phospho-mimicking mutants favor aberrant aggregation (thick arrow) over LLPS (thin arrow), whereas the arginine methylation-mimic favors LLPS (thick arrow) over aberrant aggregation (thin arrow). This panel was made with BioRender.

Given these outcomes, we next compared the effects of phosphorylation and methylation mimics on TDP-43 aggregation propensity using an *in vitro* aggregation assay. Recombinant TDP-43-MBP protein constructs were incubated with TEV protease and their aggregation was monitored over time. Under these conditions, selective cleavage of the MBP tag by TEV protease results in the formation of solid-phase TDP-43 aggregates and fibrils (Cook et al., 2020), indicated by an increase in turbidity. Here, we found that there were only minor differences in aggregation between TDP-43^WT^ and the phosphomimics TDP-43^S292E^ and TDP-43^S409E:S410E^ (Figure 5D). By contrast, the phosphomimic TDP-43^S292E:S409E:S410E^ and the arginine methylation mimic, TDP-43^R293F^, exhibited modestly reduced aggregation (Figure 5D). However, compared to TDP-43^WT^ this reduced aggregation was only significant for TDP-43^R293F^ (Figure 5E). Strikingly, plotting the normalized aggregation against the LLPS propensities of TDP-43^WT^ and its mutant forms, we found that the phosphomimetic mutants have a relatively higher tendency for aggregation over LLPS (Figure 5F). By contrast, TDP-43^R293F^ has a relatively higher tendency to undergo LLPS over aggregation (Figure 5F). Taken together, our results imply that TDP-43 phosphorylation and methylation may have opposing effects on TDP-43 (Figure 5F). Phosphorylation at S292, S409, and S410 reduces the propensity for TDP-43 to undergo LLPS, but has limited effects on TDP-43 aggregation (Figure 5G). Thus, S292, S409, and S410 phosphorylation may divert TDP-43 toward aggregation and away from LLPS (Figure 5G). This finding could explain why phosphorylated TDP-43 accumulates as aggregates in the urea fraction in cells. In contrast, R293 methylation allows TDP-43 to undergo normal LLPS, but reduces TDP-43 aggregation (Figure 5G). Thus, R293 methylation may reduce the propensity of TDP-43 to aggregate and enter the urea fraction in cells.

### Arginine methylation regulates TDP-43 aggregation in human neuronal cells

We next investigated the impact of arginine methylation on TDP-43 in a cellular context. SH-SY5Y cells were treated with AdOx, an arginine methyltransferase inhibitor, followed by fractionation and western blot analysis. We found that global methyltransferase inhibition promoted the accumulation of TDP-43 in the urea fraction (Figure 6A and B). Conversely, the overexpression of PRMT1 led to a decrease in TDP-43 aggregation, evident by a decrease of total and phosphorylated TDP-43 in the urea fraction (Figure 6C and D). To further investigate the effect of hypomethylation on TDP-43 aggregation, we next mutated TDP-43 to have an additional RGG motif (TDP-43^G308R^) (Figure 4A). Based on previous studies, the introduction of an additional RGG motif has led to the hypomethylation and subsequent aggregation of another ALS-linked RBP, FUS (Qamar et al., 2018). Accordingly, we found that TDP-43^G308R^ significantly accumulated in the urea fraction and was more highly phosphorylated at S409/S410, compared to TDP-43^WT^ (Figure 6E and F). Together, these findings suggest that arginine methylation exerts a protective role on TDP-43, perhaps by decreasing its aggregation propensity via regulation of phosphorylation.

**Figure 6.**
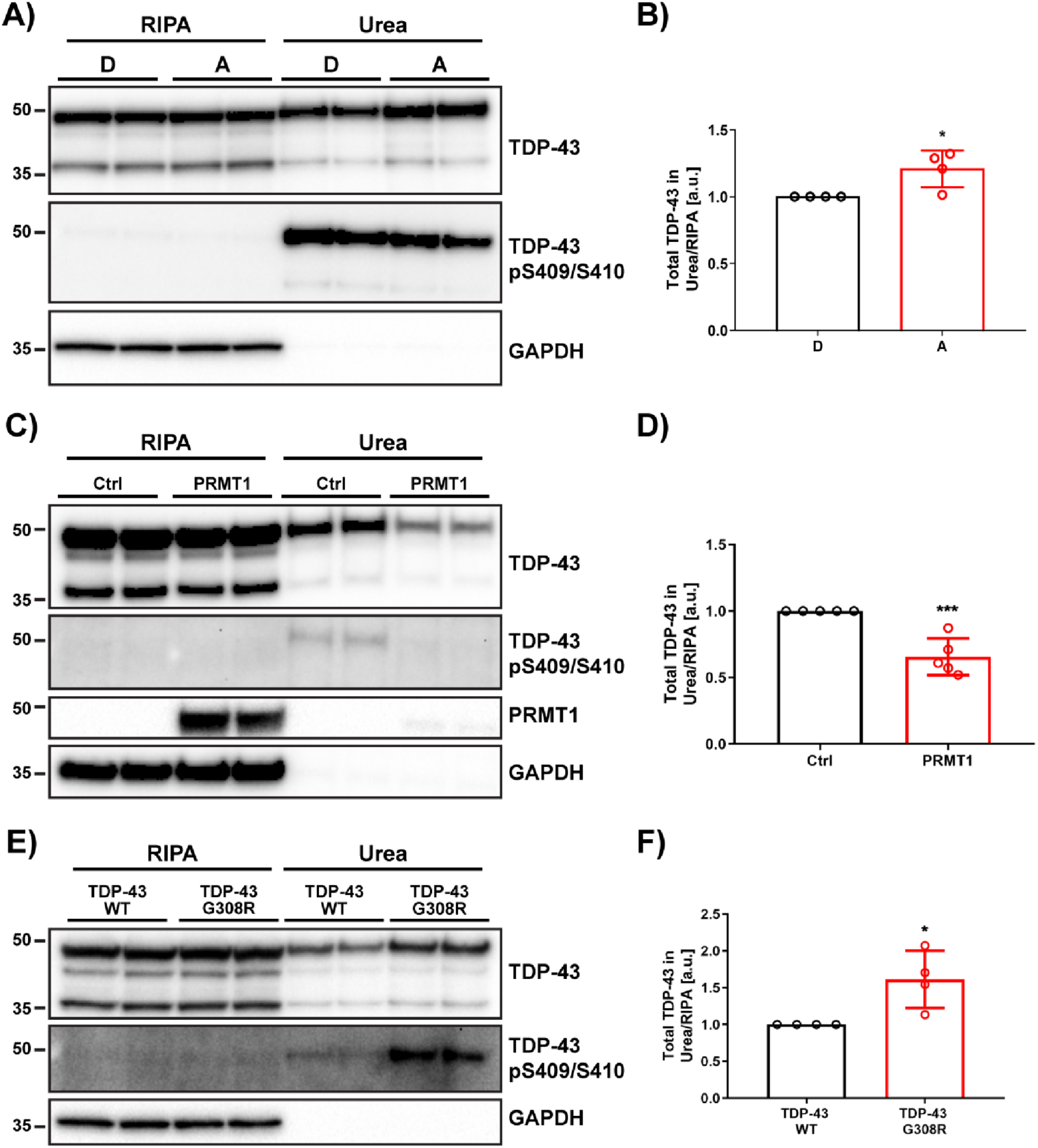
Methylation of TDP-43 regulates its aggregation. **A)** Western blot of total and pTDP-43 in RIPA and urea fractions of TDP-43^WT^-transfected SH-SY5Y-cells treated with DMSO (D) or with methyltransferase inhibitor AdOx (A) at a final concentration of 20 µM for 24h. **B)** Quantification of urea/RIPA ratio of total TDP-43 normalized to levels in DMSO-treated cells (mean band signal ± SD, unpaired t test, n = 4). **C)** Western blot of total and pTDP-43 in RIPA and urea fractions of TDP-43^WT^-transfected SH-SY5Y-cells co-transfected with empty control plasmid (Ctrl) or PRMT1. **D)** Quantification of urea/RIPA ratio of total TDP-43 normalized to levels in Ctrl-plasmid co-transfected cells (mean band signal ± SD, unpaired t test, n = 5). **E)** Western blot of total and pTDP-43 in RIPA and urea fractions of TDP-43^WT^ or TDP-43^G308R^ transfected SH-SY5Y-cells at time point of 24h post-transfection. **F)** Quantification of urea/RIPA ratio of total TDP-43 normalized to levels in TDP-43^WT^-transfected cells (mean band signal ± SD, one-way ANOVA with Dunnett’s multiple comparison test, n = 4). ^∗^p < 0.05, ^∗∗∗^p < 0.001.

### Crosstalk between TDP-43 arginine methylation and p38**α**-mediated phosphorylation

The observation that TDP-43 undergoes PRMT1-mediated arginine methylation at the R293 site, coupled with our purified protein data suggesting contrasting outcomes between TDP-43 phosphorylation and methylation, led us to ask whether phosphorylation at S292 interferes with arginine methylation at the adjacent residue, R293, or vice versa (Figure 4A). To answer this question, we expressed flag-tagged TDP-43^WT^ as well as TDP-43^S292E^, TDP-43^S292N^, and TDP-43^S292A^ mutants in SH-SY5Y cells and analyzed their methylation status. Immunoprecipitation followed by immunoblotting revealed a striking reduction in the MMA-signal in the phospho-mimicking TDP-43^S292E^ mutant (Figure 7A). Interestingly, when compared to TDP-43^WT^, the ALS-linked TDP-43^S292N^ mutant showed a modest decrease in MMA-levels, whereas the TDP-43^S292A^ mutant did not show any difference (Figure 7A). As shown previously (Figure 3C), the TDP-43^S292E^ mutant also promoted phosphorylation of TDP-43 at S409/S410, further underlining the anti-correlative relationship between TDP-43 arginine methylation and S409/S410 phosphorylation (Figure 7A). These observations suggest that phosphorylation at S292, possibly combined with increased phosphorylation at S409/S410, could interfere with TDP-43 methylation. However, these findings do not rule out that a reduction in TDP-43 methylation could be due to steric hindrance caused by the glutamic acid residue at position 292.

**Figure 7.**
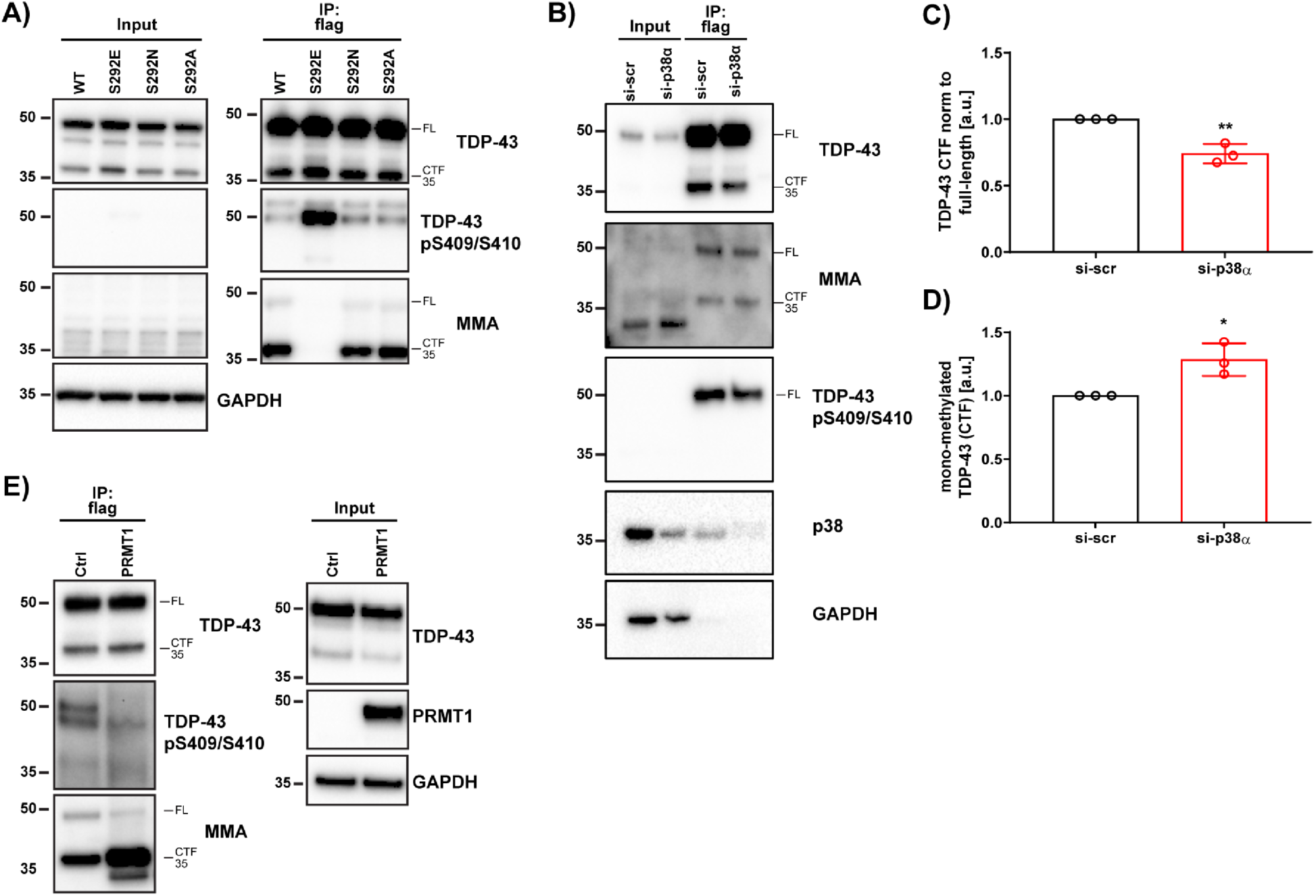
Crosstalk between PRMT1-catalyzed arginine methylation and p38α-mediated phosphorylation of TDP-43. **A)** Western blot of immunoprecipitated flag-tagged TDP-43 from SH-SY5Y cells probed with antibodies against TDP-43, pTDP-43 and MMA. GAPDH serves as a loading control. **B)** Western blot of immunoprecipitated flag-tagged TDP-43^WT^ from the SH-SY5Y cells with and without siRNA-induced p38α knockdown probed with antibodies against TDP-43, p38α, pTDP-43 and MMA. GAPDH serves as a loading control. **C)** Quantification of TDP-43-CTF/full length-ratio (mean band signal ± SD, unpaired t test, n = 3). (^∗^p < 0.05, ^∗∗^p < 0.01). **D)** Quantification of mono-methylated TDP-43-CTF normalized to total TDP-43-CTF (mean band signal ± SD, unpaired t test, n = 3). (^∗^p < 0.05, ^∗∗^p < 0.01). **E)** Western blot of immunoprecipitated flag-tagged TDP-43^WT^ from the SH-SY5Y cells with and without PRMT1 overexpression probed with antibodies against TDP-43, pTDP-43 and MMA. GAPDH serves as a loading control.

To further define the relationship between these two PTMs, we studied the methylation status of TDP-43 after genetic depletion of p38α. As shown previously (Figure 1B), we found that p38α downregulation reduced the phosphorylation of TDP-43^WT^ at S409/S410 (Figure 7B). Interestingly, western blot analysis also revealed that p38α depletion significantly decreased the formation of the ∼35 kDa TDP-43-CTF (Figure 7B and C), and resulted in elevated levels of mono-methylated TDP-43-CTF (Figure 7B and D). These observations suggest that there is an interplay between arginine methylation and p38α-mediated phosphorylation, and that reduced p38α activity could increase TDP-43 arginine methylation. Indeed, we found that overexpression of PRMT1 led to a strong increase in mono-methylated TDP-43-CTF and a striking reduction in TDP-43 phosphorylation at S409/S410 (Figure 7E), further corroborating the hypothesis that there is crosstalk between TDP-43 arginine methylation and phosphorylation at disease-relevant residues.

## Discussion

Since the discovery that ∼97% of ALS cases and ∼50% of FTD cases present with TDP-43 proteinopathy (Arai et al., 2006; Harrison and Shorter, 2017; Neumann et al., 2006), TDP-43 has been subject to much investigation. However, the mechanisms leading to an accumulation of insoluble TDP-43 aggregates are not yet fully understood. Aberrant TDP-43 phosphorylation is one of the major distinguishing pathological features of TDP-43 inclusions in human brains (Hasegawa et al., 2008; Neumann et al., 2009). Although the consequences of these phosphorylation events have not been unequivocally established, aberrant phosphorylation of TDP-43 is associated with cytoplasmic mislocalization, decreased solubility, aberrant cleavage and cytotoxicity (Barmada et al., 2010; Brady et al., 2011; Kim et al., 2015; Liachko et al., 2010; Nonaka et al., 2009; Zhang et al., 2010). To date, casein kinases CK1 and CK2, CDC7 and TTBK1/2 have been shown to phosphorylate TDP-43 *in vitro* and *in vivo,* and promote its pathological aggregation and neurotoxicity (Carlomagno et al., 2014; Choksi et al., 2014; Goh et al., 2018; Kametani et al., 2009; Liachko et al., 2013, 2014; Liu et al., 2015; Meyerowitz et al., 2011; Nonaka et al., 2016; Sreedharan et al., 2015; Taylor et al., 2018). However, some studies have found that hyperphosphorylation of the TDP-43 PrLD can reduce TDP-43 LLPS and aggregation (Li et al., 2011; Silva et al., 2021), indicating a complex interplay between combinatorial TDP-43 PTMs and pathology. Importantly, increased activation of the MAP kinase p38 has been detected in human post-mortem ALS tissue, which is further substantiated by the findings that persistent activation of the p38 signaling pathways induce neurodegeneration (Bendotti et al., 2004; Dewil et al., 2007; Tortarolo et al., 2003). However, until now, the effect of p38*α* on TDP-43 specifically has not been explored.

In this study, we elucidated the impact of p38α on TDP-43 proteinopathy. Using neuronal cells, we first showed that genetic depletion and pharmacological inhibition of p38α suppressed TDP-43 phosphorylation, aggregation, and toxicity. We further established that TDP-43 is a substrate of p38α-mediated phosphorylation. Using *in vitro* biochemical assays followed by LC-MS/MS we established that TDP-43 is directly phosphorylated by p38α at residues S292, and S409/S410. Intriguingly, mutations at S292 have been linked to pathogenicity in both sporadic and familial ALS. However, no biochemical mechanistic data have been reported on the effect of this mutation (Xiong et al., 2010; Zou et al., 2012). Here, we demonstrated that the phospho-mimetic mutant TDP-43^S292E^ significantly promoted phosphorylation of TDP-43 at S409/S410 and enhanced the accumulation of insoluble TDP-43 aggregates. These findings identified S292 as a major site for TDP-43 phospho-regulation. Interestingly, we found that S292, and the RGG-motif immediately following this residue are highly conserved across the phylogenic spectrum. The RGG-motif presents a major site for methylation by members of the PRMT family (Chang et al., 2011; Huang et al., 2018; Thandapani et al., 2013; Wall and Lewis, 2017), and we show here that R293 is a major site for methylation by PRMT1 in human neuronal cells. This result supports earlier proteomic studies of mouse embryonic and brain tissue indicating that TDP-43 can be methylated at R293 (Guo et al., 2014; Larsen et al., 2016). Furthermore, recent evidence also corroborates the possible role of arginine methylation in neurodegeneration. Specifically, PRMT1 was identified as a significant modulator of toxicity in C9-ALS (Ortega et al., 2020).

There is increasing evidence suggesting that phosphorylation and arginine methylation co-exist on the same protein, and that these PTMs can have opposing or potentiating effects on protein function (Basso and Pennuto, 2015; Lu et al., 2017). For example, the functions of p16 protein are regulated by the antagonistic crosstalk between arginine methylation at residue R138 and phosphorylation at residue S140 (Lu et al., 2017). Here, we provide similar evidence for the crosstalk between PRMT1-catalyzed arginine methylation at R293 and p38α-mediated phosphorylation at the adjacent residue S292 in the regulation of TDP-43 LLPS and aggregation. Our *in vitro* and *in vivo* data elucidate a dichotomous relationship between methylation and phosphorylation of TDP-43. Perhaps as a protective mechanism, TDP-43 methylation at R293 reduces phosphorylation at S292 and S409/S410, which in turn reduces TDP-43 aggregation. Conversely, the phosphomimetic S292E reduces the levels of TDP-43 methylation. Our biochemical studies suggest that S292 and S409/S410 phosphorylation render TDP-43 less prone to undergo LLPS, which may divert TDP-43 along pathological aggregation trajectories (Conicella et al., 2016). Whether TDP-43 methylation at R293 inhibits phosphorylation at S292, or vice versa, due to steric hindrance have yet to be further investigated. Nevertheless, our study suggests for the first time an intricate interplay between protein phosphorylation and arginine methylation in the regulation of TDP-43. While additional studies will be necessary to further clarify the dynamics of these regulatory processes, especially in the context of more complex, clinically-relevant models, our study provides a platform for developing novel therapeutic strategies to inhibit p38α or promote PRMT1 activity to rescue TDP-43 pathologies associated with ALS/FTD. Indeed, it is interesting to note that the brain-penetrant p38*α* inhibitor, VX-745, which mitigates TDP-43 toxicity in primary neurons (Figure 1J), has reached phase 2 clinical trials for Alzheimer’s disease, Huntington’s disease, and dementia with Lewy bodies (Germann and Alam, 2020; Prins et al., 2021). Our data indicate that VX-745 might also be considered as a clinical candidate for ALS/FTD that presents with TDP-43 proteinopathy. Indeed, we suggest that strategies to reduce p38α-mediated TDP-43 phosphorylation and promote R293 methylation could have therapeutic utility for ALS/FTD and other TDP-43 proteinopathies.

## Acknowledgements

We thank Charlotte Fare and Katie Copley for feedback on the manuscript, and Rebecca Jarvis for small-molecule curation. MA and HMO were supported by the AstraZeneca postdoctoral fellowships. BC was a member of the AstraZeneca graduate program. KLM was supported by supported by a NSF graduate research fellowship (DGE-1321851). AFF was supported by NIH grants T32AG00255 and F31NS087676. EMB was supported by a Milton Safenowitz Post-Doctoral Fellowship from the ALS Association and NIH grant F32NS108598. RRC was supported by NIH grants T32GM008275, F31AG060672, and a Blavatnik Family Fellowship in Biomedical Research. JS was supported by Target ALS, The Association for Frontotemporal Degeneration, The Packard Foundation for ALS research, The ALS Association, The G. Harold and Leila Y. Mathers Charitable Foundation, the Office of the Assistant Secretary of Defense for Health Affairs (USA), through the Amyotrophic Lateral Sclerosis Research Program (W81XWH-20-1-0242), and NIH grants R01GM099836 and R21AG065854. This work was supported by a grant to NJB, DGB, JS, ADG, SF from the Target ALS Foundation and ALS Finding a Cure.

---

**Figure S1.**
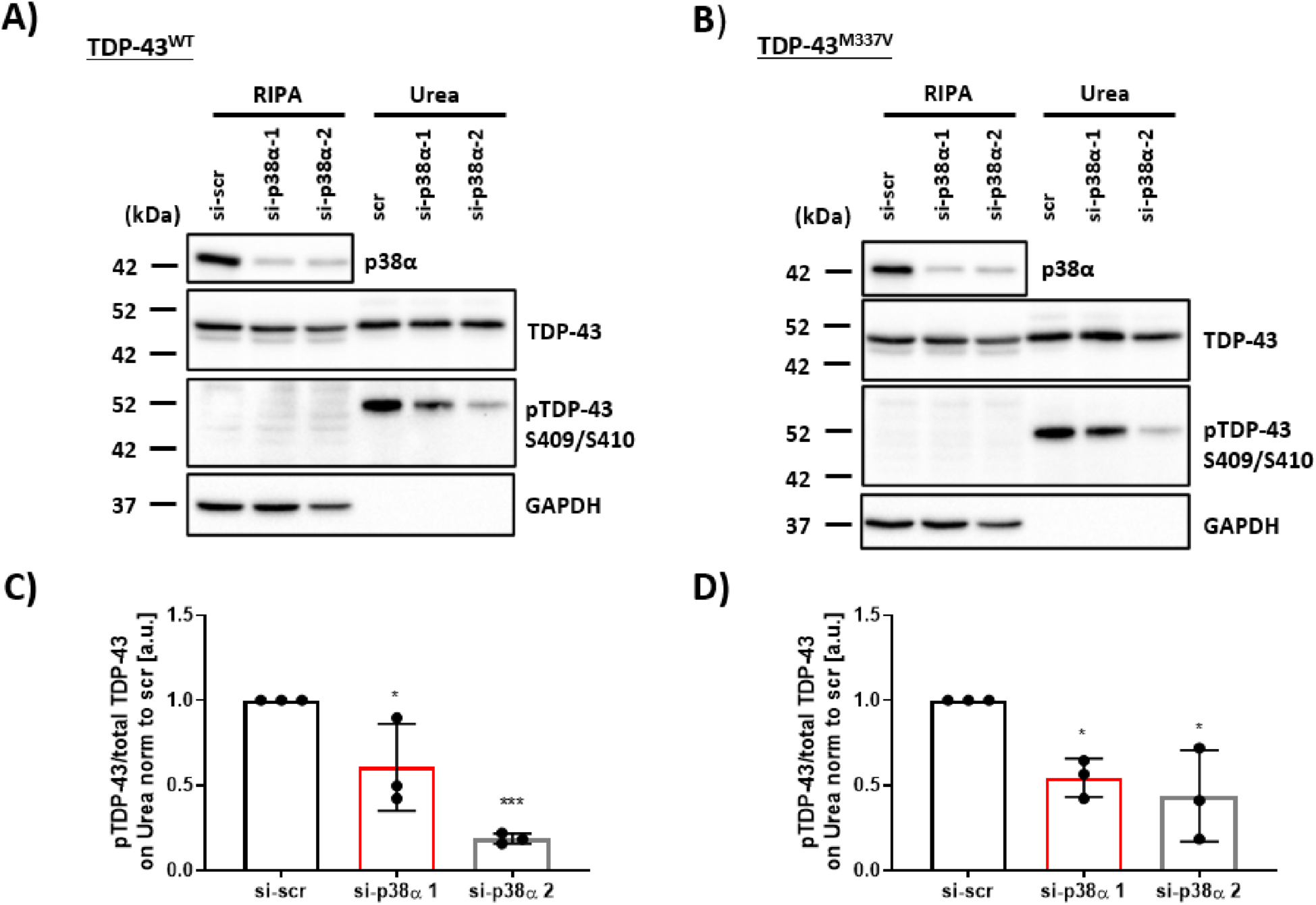
Genetic ablation of p38α MAPK reduces the aggregation and phosphorylation at S409/S410 for both TDP-43^WT^ and TDP-43^M337V^. **A)** Western blot of total and pTDP-43^WT^ in RIPA and urea fractions of cells with siRNA-induced p38α knockdown in SH-SY5Y cells after 48h. GAPDH was used as a loading control. **B)** Western blot of total and pTDP-43^M337V^ in RIPA and urea fractions of cells with siRNA-induced p38α knockdown in SH-SY5Y cells after 48h. GAPDH was used as a loading control. **C)** Quantification of pTDP-43^WT^/total TDP-43^WT^ -ratio in urea fraction normalized to levels in scrambled siRNA (si-scr). **D)** Quantification of pTDP-43^M337V^/total TDP-43^M337V^ -ratio in urea fraction normalized to levels in si-scr (mean band signal ± SD, one-way ANOVA with Sidak’s multiple comparison test, n = 3). ^∗^p < 0.05, ^∗∗∗^p < 0.001. Related to Figure 1.

**Figure S2.**
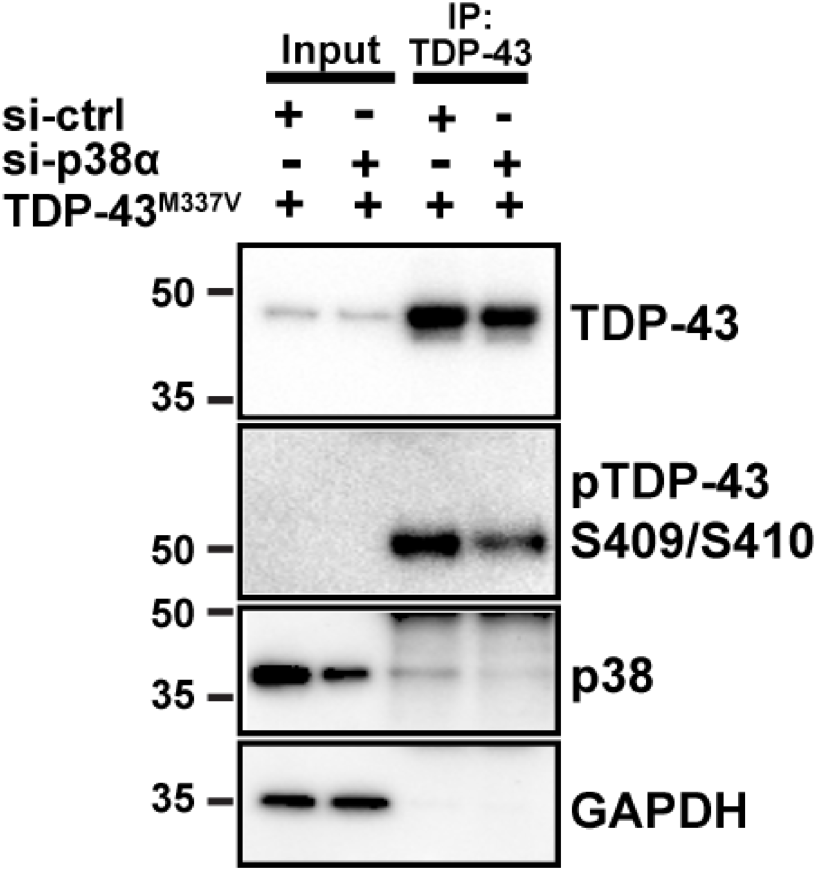
Knockdown of p38α reduces the co-immunoprecipitation of endogenous p38α with TDP-43. Western blot of immunoprecipitated TDP-43^M337V^ and co-immunoprecipitated endogenous p38α from the SH-SY5Y cells with and without siRNA-induced p38α knockdown. GAPDH serves as a loading control. Related to Figure 3.

